# Scent mark signal investment predicts fight dynamics in house mice

**DOI:** 10.1101/2022.05.19.492706

**Authors:** Caitlin H. Miller, Klaudio Haxhillari, Matthew F. Hillock, Tess M. Reichard, Michael J. Sheehan

## Abstract

Signals mediate competitive interactions by allowing rival assessment, yet are often energetically expensive to produce. Individuals face tradeoffs when deciding when and where to signal, such that over or under-investing in signaling effort can be costly. One of the key mechanisms maintaining signal reliability is via social costs. While the social costs of over-signaling are well-known, the social costs of under-signaling are underexplored, particularly for dynamic signals. In this study we investigate a dynamic and olfactory-mediated signaling system that is ubiquitous among mammals: scent marking. Male house mice territorially scent mark their environment with metabolically costly urine marks. While competitive male mice are thought to deposit abundant scent marks in the environment, we recently identified a cohort of low-marking males that win fights. Whereas there are clear energetic costs to investing in urine signals in mice, we hypothesized that there may be social costs imposed on individuals who under-invest in signaling. Here we find that scent mark investment predicts fight dynamics. Despite fight outcome being unambiguous, aggressive intensity varies considerably across trials. Males that produce fewer scent marks engage in more intense fights that take longer to resolve. This effect appears to be driven by an unwillingness among losers to acquiesce to weakly signaling winners. We therefore find evidence for rival assessment of scent marks as well as social costs to under-signaling, which supports existing hypotheses for the importance of social punishment in maintaining optimal signaling equilibria. Our results further highlight the possibility of diverse signaling strategies in house mice.

## 1. Introduction

Signals of competitive ability play an important role in mediating rival assessment in aggressive contests (1–9). However, signal production is often energetically expensive, and individuals face tradeoffs when investing in signaling effort relative to other life history traits (10–12). For example, increased signal investment can result in reduced gamete production (13–15), immune deficits (16,17), and higher risks of parasitism or predation (18–22).

In addition to production tradeoffs, there are social costs to signaling either too much or too little. Individuals that “over-signal” their competitive ability receive heightened aggression from competitors (23–28). Whereas individuals that “under-signal” struggle to establish dominance relationships (26,28). Such mismatches in signaled versus actual competitive ability muddle accurate rival assessment, resulting in escalated contests (26,28). Receiver-dependent social punishment has been hypothesized as an important mechanism in maintaining optimal signaling equilibria (23). While the social costs of over-signaling (i.e. ‘bluffing’ or ‘cheating’) have been well-examined, the social costs of under-signaling are under-studied, particularly for dynamic signals.

Here, we explore a dynamic and olfactory-mediated signaling system that is central to mammalian communication: scent marking (29–31). Scent marks persist in the environment for long periods of time (32–35) and provide a record of social relationships that can be assessed by receivers (33,35–37). Scent marks have further been proposed as ‘cheat-proof’ signals of status due to the inherent metabolic and physical challenges of maintaining a scent-marked territory (35,36).

In house mice (*Mus musculus domesticus*), urine marking is arguably the most prominent signaling modality. The generally accepted canon is that competitive males are aggressive, territorial, and mark highly (38–41). In addition to the costs of actively re-marking and patrolling a territory, urine marks themselves are metabolically costly in house mice (36,42–44). Urine marking has previously been shown to have important life history costs in house mice, as males that invest in marking earlier in life experience reduced body growth (42). It is, therefore, generally assumed that urine marking is an honest indicator of a male’s status and competitive ability (38–41,45–47). Yet, we have recently tested this assumption and found it to be incomplete—urine marking prior to a contest did not predict wins or losses among size-matched rivals, in part due to the presence of low-marking competitive males (48).

This surprising result led us to ask whether and how male house mice use scent mark information in competitor assessments. The objectives of this study were to: (1) test the hypothesis that scent mark signaling prior to a fight shapes contest dynamics, and (2) examine the potential social costs of under-signaling. We predicted that high quality individuals that accurately signaled their competitive ability would beneficially engage in less intense aggressive behaviors, and more quickly resolve their fights. In contrast, individuals that under-signaled their competitive ability would face the social costs of escalated aggressive encounters, and experience delayed contest resolution.

## 2. Material & Methods

### (a) Study system

To explore scent marking and aggressive behaviors we used male house mice (n=62), as males will competitively urine mark and exhibit territorial aggression (33,39–41,46,49–51). Experimental individuals were from two wild-derived inbred strains (NY2 and NY3) of house mice (52). The progenitors of these strains were captured near Saratoga Springs, NY in 2013 by MJS and are related to the SarA/NachJ, SarB/NachJ and SarC/NachJ strains now available from the Jackson Lab. We used two wild-derived strains because competitive behaviors are less pronounced in highly inbred and domesticated laboratory strains (53,54), and individuals within inbred strains tend to share identical urinary protein profiles (55). At the time of experimentation all males were adult (3-5 months old) and sexually experienced. Mice were housed in an Animal Care facility at Cornell University with a 14:10 shifted light:dark cycle (dark cycle: 12PM–10PM), with food and water provided *ad libitum*. To reduce handling stress confounds, mice were transferred between their home cage and the experimental arena using transfer cups (56).

### (b) Scent mark signaling and aggressive contests

In our previous work examining signal allocation decisions, we were surprised to find that scent marking behavior did not clearly predict wins or losses during fights, and instead identified a cohort of low-signaling competitive males (48). Together, these results led us to investigate the aggressive contests within this dataset in greater detail to better understand the relationship between signaling and competitive ability (48).

We placed males in an arena separated by a mesh barrier where they could see, hear, and smell each other but were limited to minimal physical contact (**Figure 1A**). This allowed us to measure male urine marking prior to a contest. After 30 minutes, we removed the mesh barrier and males engaged in a fight trial for an additional 30 minutes (**Figure 1A**). Trials were performed on filter paper to prevent smearing of urine marks, for easier detection of urine deposition events. One day prior to experimentation, we recorded male body weights to size-match individuals. As house mice are nocturnal, we conducted all experiments during the dark cycle between 12 PM-5 PM. Age and weight-matched adult breeding males of distinct wild-derived strains (NY2 and NY3) were paired as competitors, resulting in a total of 31 pairs (n=62). We therefore ensured that no two paired competitors were genotypically identical and that their scent marks were perceptibly different (i.e. characterized by unique major urinary protein profiles) (44,46,55,57,58). We ear-clipped and bleached a patch of rump fur of one male in each pair a week prior to experimentation for easy identification of males within a pair (NY2 and NY3 strains are visibly indistinguishable).

**Figure 1.**
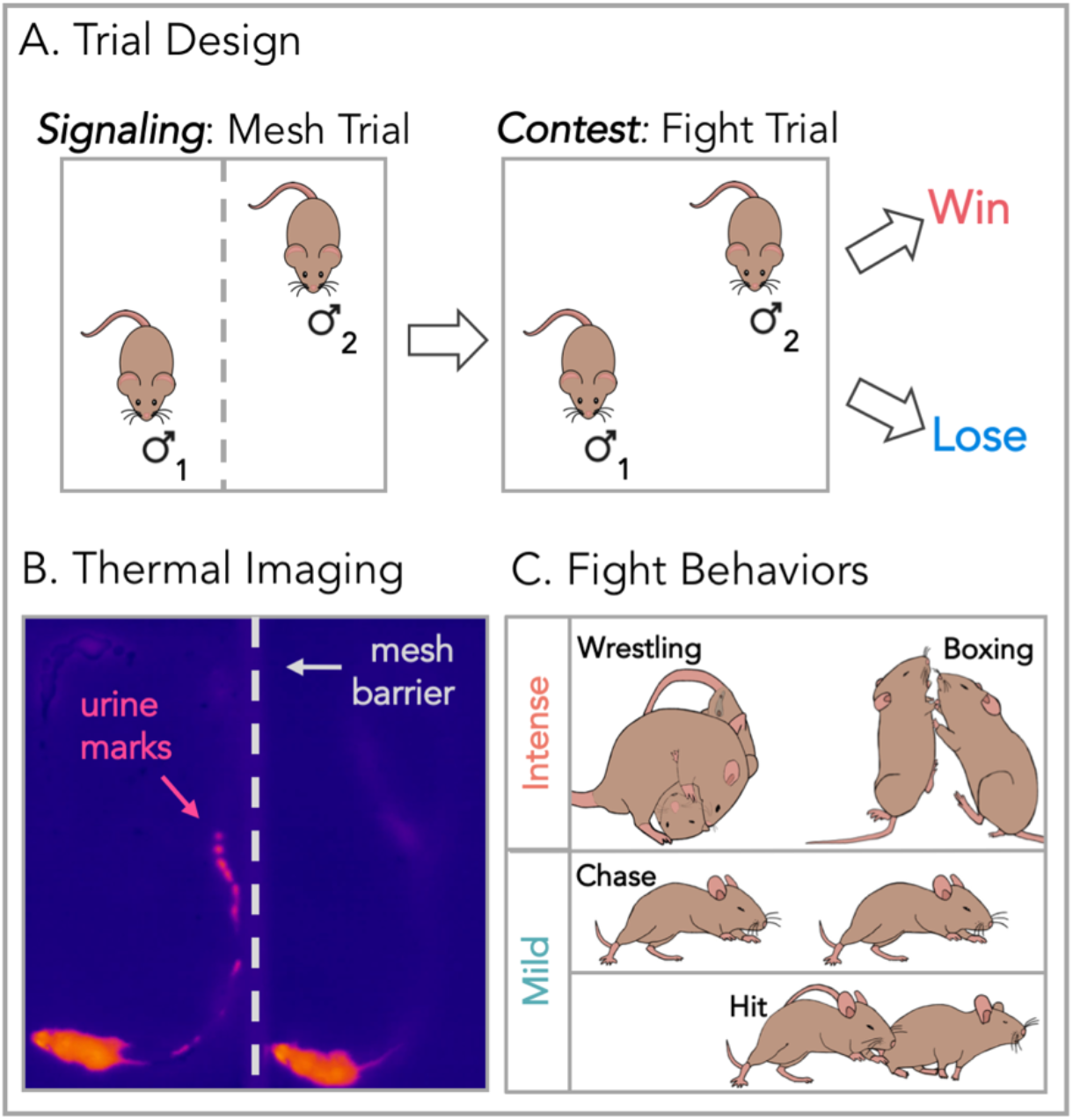
Trial design and recording methods. **(A)** Two-part trial design starting with a 30-minute signaling trial where paired competitors were separated by a mesh barrier, and urine marking was measured. The mesh barrier was removed and males entered into the contest phase of the trial (fight trial) for an additional 30 minutes. **(B)** Urine depositions were recorded using thermal imaging. Urine exits the body hot and then cools below substrate temperatures, providing a distinct thermal signature. **(C)** For each fight trial, four aggressive behaviors were scored: wrestling, boxing, chases, and hits. Wrestling and boxing were classified as intense attacks; chases and hits were classified as mild attacks.

All trials were recorded with a thermal imaging camera system (PI 640; Optris Infrared Sensing; 33° x 25° lens; ~3 Hz; thermal detection: 61°F - 107°F) and a security camera system (iDVR-PRO CMS; 1080p; 30 fps). Thermal imaging allowed for the detection of urine mark deposition events with fine spatiotemporal detail. Urine leaves the body hot (close to body temperature) and quickly cools to below the ambient substrate temperature, providing a distinctive thermal signature (**Figure 1B**). The security camera system was used to visualize high-speed aggressive encounters. Both systems were used to cross-check for recording errors.

Videos were scored blindly using Behavioral Observation Research Interactive Software (BORIS) (59). Urine depositions were scored as a clear hot spot in the focal mouse’s trajectory that subsequently cooled below substrate temperature (**Figure 1B**). Based on the total aggressive behaviors performed by each male, males were unambiguously classified as winners or losers (**Figures 2A & S2**). Males were further categorized as low-marking or high-marking based on whether the total urine marks deposited in the “mesh trial” (pre-fight) fell below or above the median **(Figures 1A & S1A**). These classifications were used to interrogate interactions between signal investment and fight dynamics. The following behaviors were scored in “fight trials”: chasing, hitting, boxing, and wrestling bouts (60–62) based on which male initiated these behaviors (**Figure 1C & S2A**). Aggressive behaviors were further categorized as mild or intense based on the risk of injury (i.e. belly exposure and likelihood of bites occurring) and the extent of physical contact. Chases and hits were classified as mild attacks, while boxing and wrestling bouts were classified as intense attacks (**Figure 1C**). Importantly, intense attacks are interactive behaviors, which require that the male receiving the attack actively defends themselves rather than fleeing from the interaction. No mice experienced sustained injury in these trials.

**Figure 2.**
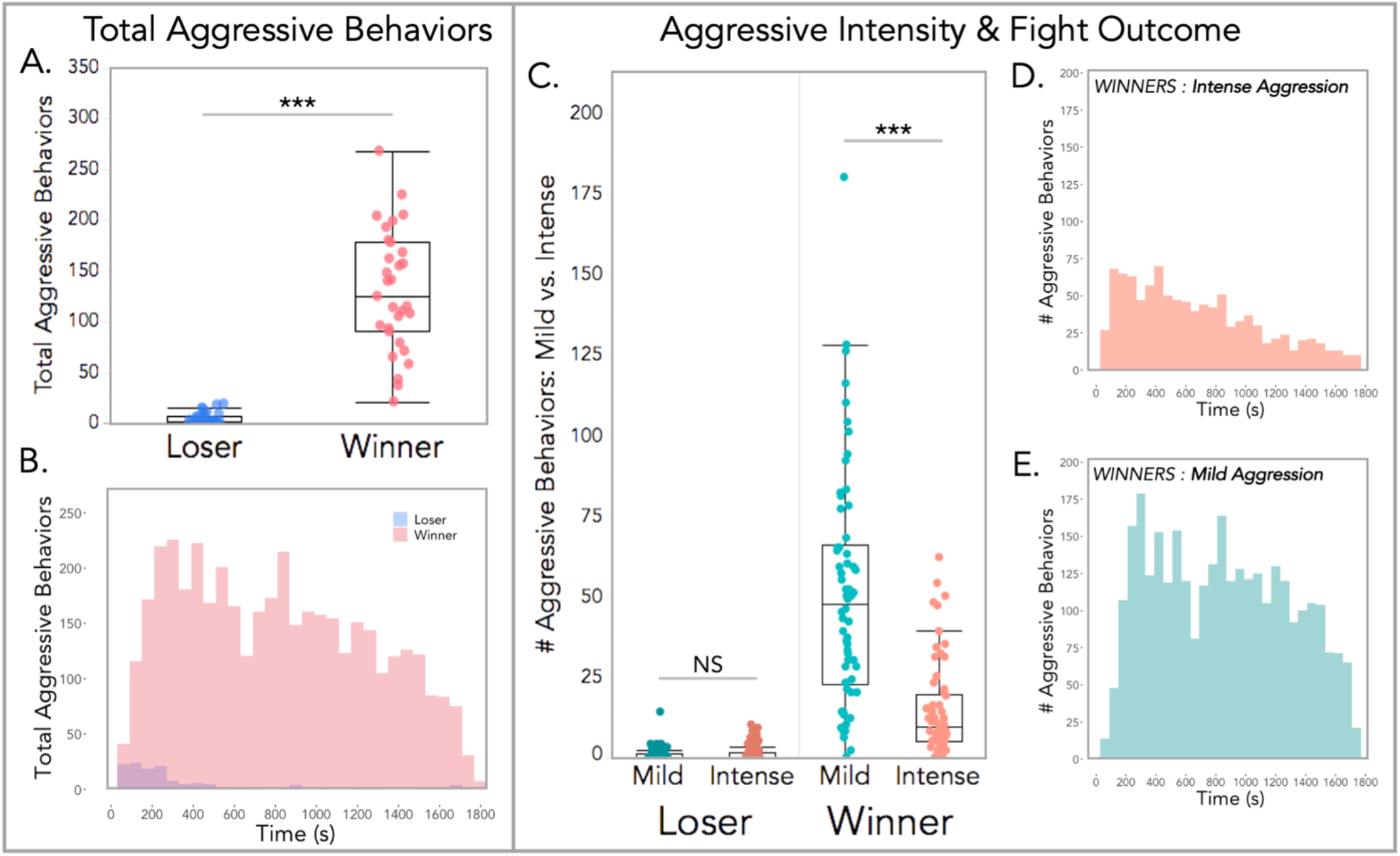
Winners displayed more aggressive behaviors throughout the fight trial, while losers rarely displayed any aggression after the first five minutes. **(A)** Total aggressive behaviors performed by males that either won or lost the fight. **(B)** Histogram of the temporal distribution of aggressive behaviors performed by winners and losers over the fight trial duration. **(C)** Total mild vs. intense aggression displayed by winners and losers. **(D,E)** Histograms of intense vs. mild aggression exhibited by winning males over the course of the fight trial. **(A,C)** Linear mixed models were used to model relationships (**Table S1)**. Analyses of variance were used to test for overall effects. Dependent variables (# aggressive behaviors) were logarithmically transformed to meet assumptions for model residuals. Significance codes: NS p>0.05, * p<0.05; ** p<0.01, *** p<0.001.

### (c) Statistical analyses

We conducted all statistical analyses in R 3.6.0 (R Development Core Team 2019). We used linear mixed models and paired statistical tests to examine relationships between dependent and response variables (**Tables S1–S4**). Models were fitted using the package *lme4* (63). The *lmerTest* package was used to calculate degrees of freedom (Satterthwaite’s method) and p-values (64). Dependent variables were logarithmically transformed for a subset of models to meet assumptions for model residuals (**Tables S1–S4**). We used a type 3 analysis of variance to test for overall effects of fixed factors or interactions in the models. Post hoc comparisons were conducted using the *emmeans* package (65). R script and data sheets used for all statistical analyses are provided (see Data Availability).

## 3. Results

### (a) Contest outcomes and aggressive intensity

Fight outcome was unambiguous in all contest pairings (**Figure S2**). Winners performed significantly more aggressive behaviors than losers (M1: *F*_1,60_= 287, *p* < 0.0001; **Figure 2A & Table S1**). This was true for both cumulative aggression (**Figure 2A**), as well as for specific fight behaviors (**Figure S2B**). Across all fight trials, winners performed 21-268 total attacks, while losers performed 0-19 (**Figure S2C**). Within pairs, the difference in attack count ranged from 21-266, with an average attack difference of 126+/-11 between competitors. These winner-loser relationships were typically apparent within the first 5 minutes, as losing males quickly halt aggression (**Figure 2B**). Winners on the other hand, rapidly escalated aggression with peak activity occurring at ~300 seconds, followed by a gradual decline (**Figure 2B**). Males thus performed fast competitor assessments once they physically engaged. While males were weight-matched as closely as possible, some variation in body weight was inevitable (**Figure S3A**). Body weight moderately predicted the total aggressive behaviors performed by individuals (M2: *F*_1,59_ = 4.5, *p* = 0.04; **Table S1)**, however including body weight resulted in a worse model overall (M1 vs. M2: **Table S1**). Body weight also did not differ between winning and losing males across trials (*t*_1,60_ = 0.4, *p* = 0.69; **Figure S3B**).

We further explored the intensity of aggressive behaviors initiated by competitors during contests and found that aggressive intensity has a significant interaction with fight outcome (M3: *F*_1,184_ = 64, *p* < 0.0001; **Figure 2C & Table S1**). Losers exhibit similarly few mild and intense aggressive behaviors (M3: *t*_1,184_ = 1.5, *p* = 0.39; **Figure 2C & Table S1**). In contrast, winners perform significantly more mild attacks than intense ones (M3: *t*_1,184_ = −9.8, *p* < 0.0001; **Figure 2C & Table S1**). Given the low rates of aggressive attacks performed by the eventual contest losers, we focused on the dynamics of aggressive behaviors initiated by the ultimate contest winners. Among winners, the temporal dynamics reveal the number of intense attacks steadily declines over the course of the fight (**Figure 2D**). Whereas mild attacks remain elevated for longer and decline in frequency more slowly (**Figure 2E**). These data indicate that while fight outcome is straightforward, attack frequency varies with intensity. Furthermore, there appear to be distinct temporal patterns for mild and intense fight behaviors.

### (b) Scent mark signaling prior to a contest predicts fight dynamics

We next explored the relationship between scent mark signaling and fight dynamics among winning males, as these individuals initiated the vast majority of aggressive behaviors (**Figure 2**). Males were categorized as either low or high-marking individuals, based on whether their total number of urine deposition events fell below or above the median number of marks (**Figures 3A & S1A**). This categorization is supported by our prior work, which shows that initial mark investment predicts marking levels days later (**Figure S1**; 48). In other words, males that are low or high-marking adhere to their respective marking groups days after an aggressive contest (48).

**Figure 3.**
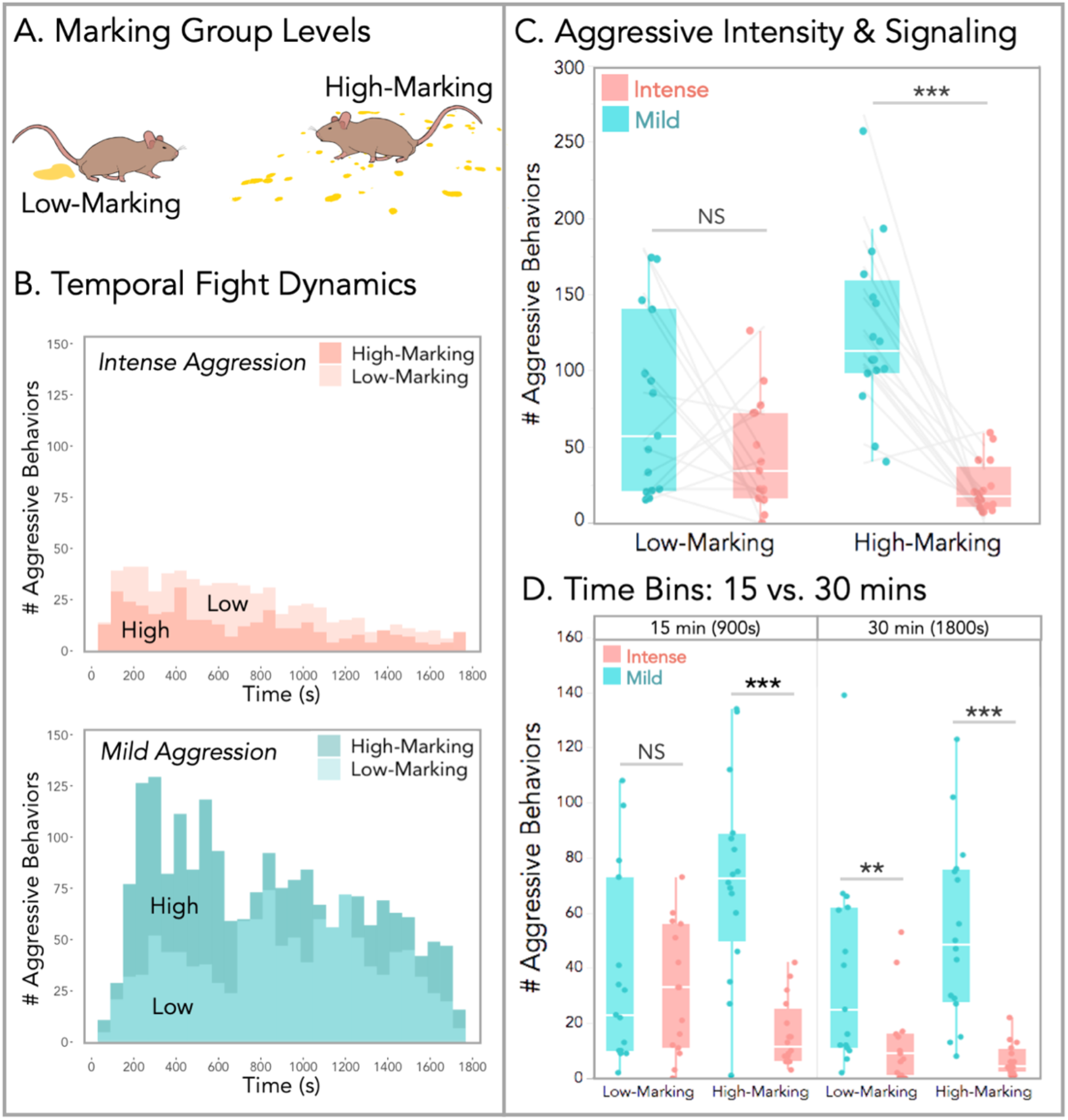
Temporal fight dynamics and intensity vary with initial signaling effort of winning males. **(A)** Males were categorized as low or high-marking individuals based on whether the total number of marks deposited prior to the fight trial fell either below or above the median (**Figure S1A**). **(B)** Histograms of the temporal distributions for intense (top) and mild (bottom) aggressive behaviors for the two marking groups (high vs. low-marking). **(C)** Boxplot of total aggressive behaviors by marking group and fight intensity. **(D)** Boxplot of total aggressive behaviors by marking group, attack intensity, and fight trial 15-minute time bins. **(C,D)** LMMs were used to model relationships, and analyses of variance were used to test for overall effects. Dependent variables were logarithmically transformed to meet assumptions for model residuals. Significance codes: NS p>0.05, * p<0.05; ** p<0.01, *** p<0.001.

The total number of attacks initiated by winners did not significantly differ between low and high-marking males (M4: *F*_1,29_ = 3.3, *p* = 0.08; **Table S2**). However, striking patterns emerged when we inspected the intensity of aggressive behaviors. We first examined the temporal distribution of mild and intense attacks for low and high-marking winners (**Figure 3B**). Low-marking winners performed more intense attacks across the fight duration (**Figure 3B**). While an inverse relationship is observed for mild aggression. High-marking winners performed dramatically more mild attacks than low-marking winners, particularly in the first 15 minutes of the fight trial (**Figure 3B**).

We further modeled the effects of fight intensity and signaling effort on contest aggression (**Figure 3C & Table S2**). The signaling effort of winners significantly predicted aggressive intensity (M5: *F*_1,58_ = 32, *p* < 0.0001; **Table S2**). Interestingly, initial marking effort did not predict aggressive behaviors in other dyadic comparisons. Initial signaling efforts of the eventual losing males did not predict loser aggression (M8: *F*_1,58_ = 0.59, *p* = 0.44; **Figure S4A & Table S4**) or winner aggression (M9: *F*_1,58_ = 1.2, *p* = 0.27; **Figure S4A & Table S4**). Similarly, the marking effort of winners did not predict loser aggression (M10: *F*_1,58_ = 1.1, *p* = 0.29; **Figure S4B & Table S4**). Moreover, body weight did not differ across low or high-marking winners and losers (**Figure S3C**).

### (c) Social costs of under-signaling

We next examined the social costs of under-signaling, and found a significant interaction between fight intensity and initial marking effort (M5: *F*_1,58_ = 6.8, *p* = 0.01; **Figure 3C & Table S2**). Winning males that invest more in scent marking perform fewer intense attacks and more mild attacks (**Figure 3C**). Whereas, males that invest less in marking engage in more intense aggression (**Figure 3C**). Comparing rates of each type of aggression per winner revealed that low-marking winners do not differ in their levels of attack intensities (M5: *t*_1,29_ = −2.1, *p* = 0.14; **Figure 3C & Table S2**). The opposite is true for high-marking winners, which perform significantly more mild relative to intense attacks (M5: *t*_1,29_ = −5.9, *p* < 0.0001; **Figure 3C & Table S2**).

Given that the differences in fight intensity toward signaling effort occurred specifically among winners, we further interrogated the temporal dynamics of these behaviors. To do this we split the fight trial into two 15-minute time bins, corresponding to the first and second half of the trial (**Figure 3D**). The differences in fight dynamics are particularly stark in the first 15 minutes (**Figure 3D**). Time bin has a strong effect on contest aggression (M6: *F*_1,87_ = 11, *p* = 0.002; **Table S2**), with a significant two-way interaction between fight intensity and time bin (M6: *F*_1,87_ = 5.9, *p* = 0.02; **Table S2**). Comparing rates of mild and intense acts of aggression per winner revealed that in the first half of the trial, low-marking winners are performing the same levels of mild and intense attacks (M6: *t*_1,87_ = −0.45, *p* = 1.0; **Figure 3D & Table S2**). While high-marking winners perform dramatically more mild attacks relative to intense attacks (M6: *t*_1,87_= −4.4, *p* = 0.0003; **Figure 3D & Table S2**). This difference in fight intensity between marking groups diminishes in the second half of the trial, such that all winners display significantly higher rates of mild compared to intense aggression (**Figure 3D**). Thus, the fights of low-signaling winners exhibit more severe escalation in the first 15 minutes, suggesting males that under-signal take longer to resolve aggressive contests than males investing more in signaling effort.

Together, these data indicate that the relative proportion of intense aggressive behaviors males perform varies with signaling effort. We assessed the proportions of specific fight behaviors executed by low and high-marking individuals (**Figure 4A**). This demonstrates that low-marking winners perform considerably more intense aggression (i.e. wrestling bouts and boxing matches; **Figure 4A**). The aggressive behaviors of high-marking individuals, however, are heavily skewed towards mild aggression (i.e. chases and hits; **Figure 4A**). We further examined the proportion of intense-to-overall attacks relative to the signaling efforts of each winning male prior to the contest (**Figure 4B**). The proportion of intense attacks is predicted by marking group (M7: *F*_1,28_ = 8.8, *p* = 0.006; **Table S3**). This illustrates a striking delineation between low and high-signaling competitive males (**Figure 4B**). Furthermore, it reveals what appear to be two distinct groups of males that each comprise a quarter of all winners: (1) intensely aggressive low-marking males and (2) mildly aggressive very high-marking males (**Figure 4B**).

**Figure 4.**
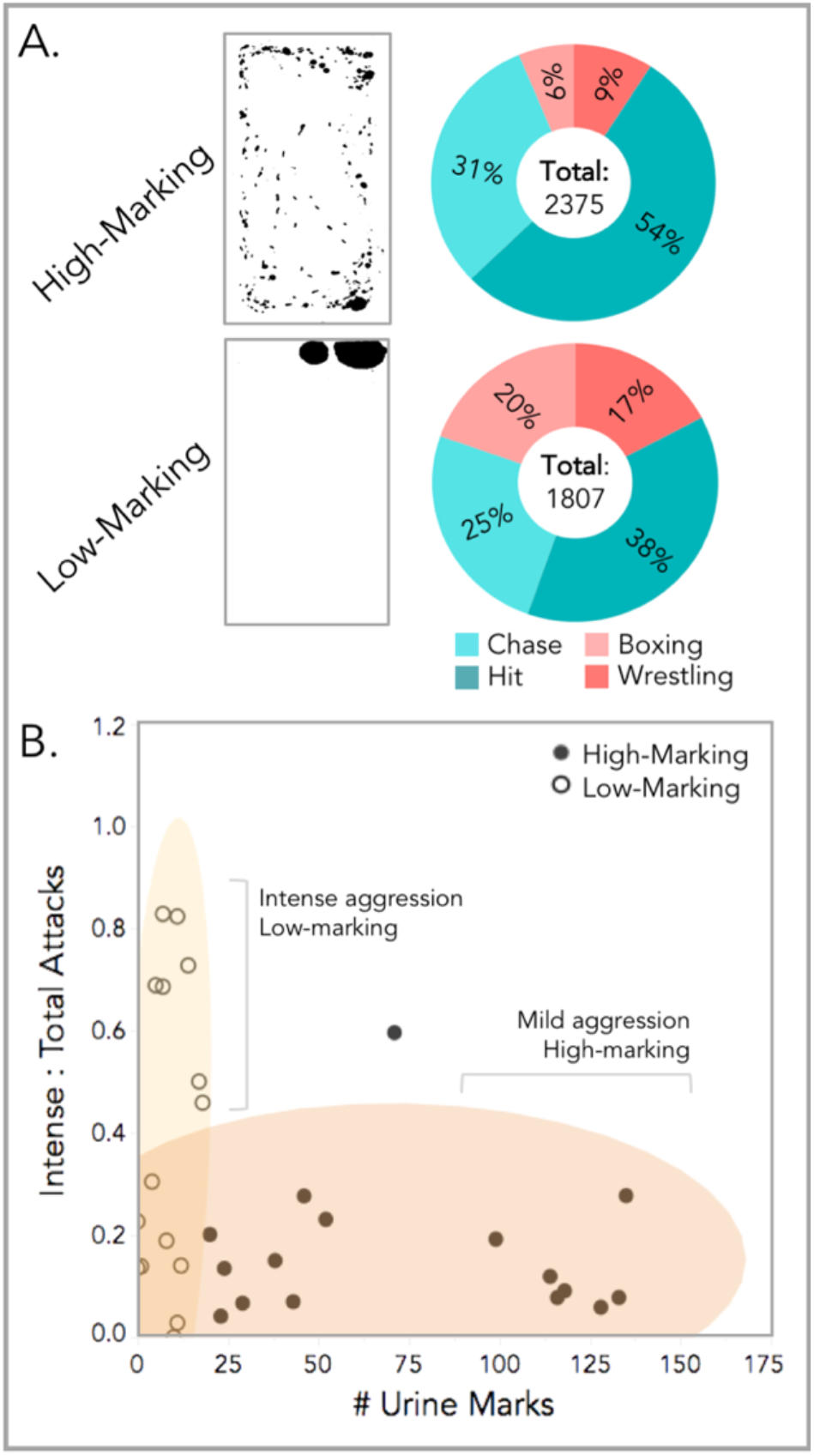
Proportions of mild and intense aggression. **(A)** Example processed urine blots of high and low-marking males (urine spots shown in black). Donut plots depict proportions of individual fight behaviors for the two marking groups **(B)** The proportion of intense : total attacks by the total number of urine marks deposited prior to the fight. Low and high-marking winners are labeled. Ellipses indicate 90% data coverage.

## 4. Discussion

Here we have shown that despite fight outcome being overwhelmingly clear (**Figure 2**), there are stark differences in contest dynamics depending on how males signaled prior to a fight. We find evidence for social costs to under-signaling in house mice, as low-marking winners experienced more intense fights and delayed contest resolution (**Figures 3 & 4**). This suggests there are likely important tradeoffs underlying signal investment decisions in terms of competitor assessment and aggression. Particularly as in our prior work, we identified a cohort of competitive yet stably low-marking male mice (48). Our work underlies the complex decisions animals face when determining their signal investment and willingness to engage in aggressive encounters. At any given moment, individuals confront metabolic resource limitations. Deciding when and where to invest these resources has important fitness consequences.

The observed differences in aggressive intensity in male house mice could be driven by winners, losers, or a combination of the two. A winner-driven explanation is that winners allocate more effort toward aggression rather than signaling, such that the total energy invested is constant across low-marking and high-marking winners. Alternatively, losing males may be less inclined to back down during attacks initiated by weakly signaling males. Importantly, intense aggressive behaviors (i.e. wrestling and boxing) are highly interactive, and require that losers actively defend themselves rather than flee from the encounter (i.e. chases). This lends support for the observed differences being loser-driven, and alludes to a possible key difference between losing an encounter and submitting to a competitor.

The trial design limits males’ ability to escape the interaction. In naturalistic environments males would likely avoid prolonged encounters, and dominance relationships would be established through shorter repeated interactions. Nevertheless, the sustained encounters used in this study allowed us to observe temporal shifts in fight intensity. We find that low-signaling winners don’t transition to more mild aggression until midway through the fight, whereas high-signaling winners start off relatively mild. Fight resolution is therefore delayed when winners signal lowly prior to a fight. Similar to what has been observed in aggressive contests between paper wasps (26) and chameleons (28), we find that male house mice experience social costs as a result of inaccurately signaling their competitive ability. This is striking because the costs of under-signaling appear quite high. “Scent-silent” males engage in more intense fights, take longer to resolve dominance relationships, and incur greater risk of injury.

Given the potential social costs, the existence of competitive low-signaling males suggests there may be fitness benefits to remaining scent-silent, at least under some socioecological conditions. Perhaps the most obvious benefit to reduced signaling is that it may save energy. Males might withhold signal investment to build up their metabolic reserves, as urine marking is energetically expensive (42,43,47,66). In doing so, males may gain body mass and more effectively defend territories later in the season. This is plausible given that prior work has shown males who invest early in urine marking pay the cost of reduced body size (42). The low-signaling effort observed among competitive males could therefore reflect important features of life history in house mice, and potentially in many other species.

Males entering into the trials had no prior competitive experience. Males may strategically hold off scent mark investment until there are rival males present, suggesting population density may have large effects on signaling strategies (67–69). This is particularly intriguing given prior hypotheses of urine marking as ‘cheat-proof’ (35,36). These hypotheses emphasize the inability to deceptively over-report (i.e. bluff) one’s competitive ability but do not address the possibility of males under-reporting. Our results highlight the importance of investigating under-signaling strategies, as they may be more common than previously appreciated across taxa.

Another possibility is that male mice exhibit a spectrum of signaling strategies, including the classically described “territorial males” that invest highly in marking as well as scent-silent “satellite” males. In this scenario, low-signaling individuals might avoid detection by other males yet are competitive enough to mate, though reduced marking effort likely decreases the chances of obtaining mating opportunities (33,70,71). Previous work described males reducing their scent marking after losing (41). Indeed, in the trials reported here, we found that high-marking males dramatically reduce their marking efforts after losing (48). However, this scenario does not readily explain the observed patterns among winning males that continue to mark infrequently. It may be that in time they would increase investment in scent marking, and shifting to high-marking is a slower process than downregulating after a loss (43,47). Studies of scent marking effort in more natural contexts are sorely needed.

We find evidence for direct social costs as a result of under-signaling one’s competitive ability. This supports existing theoretical frameworks underlying the importance of social punishment in shaping patterns of signal investment (23,25,26). Our results further provide evidence that urine marking is used in competitor assessments and appears to determine losers’ willingness to submit. Furthermore, our findings highlight the possibility of diverse signaling strategies in house mice. As a dynamic signaling system, individuals may flexibly adjust scent mark investment depending on the social landscape and their energetic reserves. Diverse strategies may be more commonplace for dynamic signals across taxa than is currently recognized, and warrants further investigation.

## Supplemental Materials

**Table S1.**
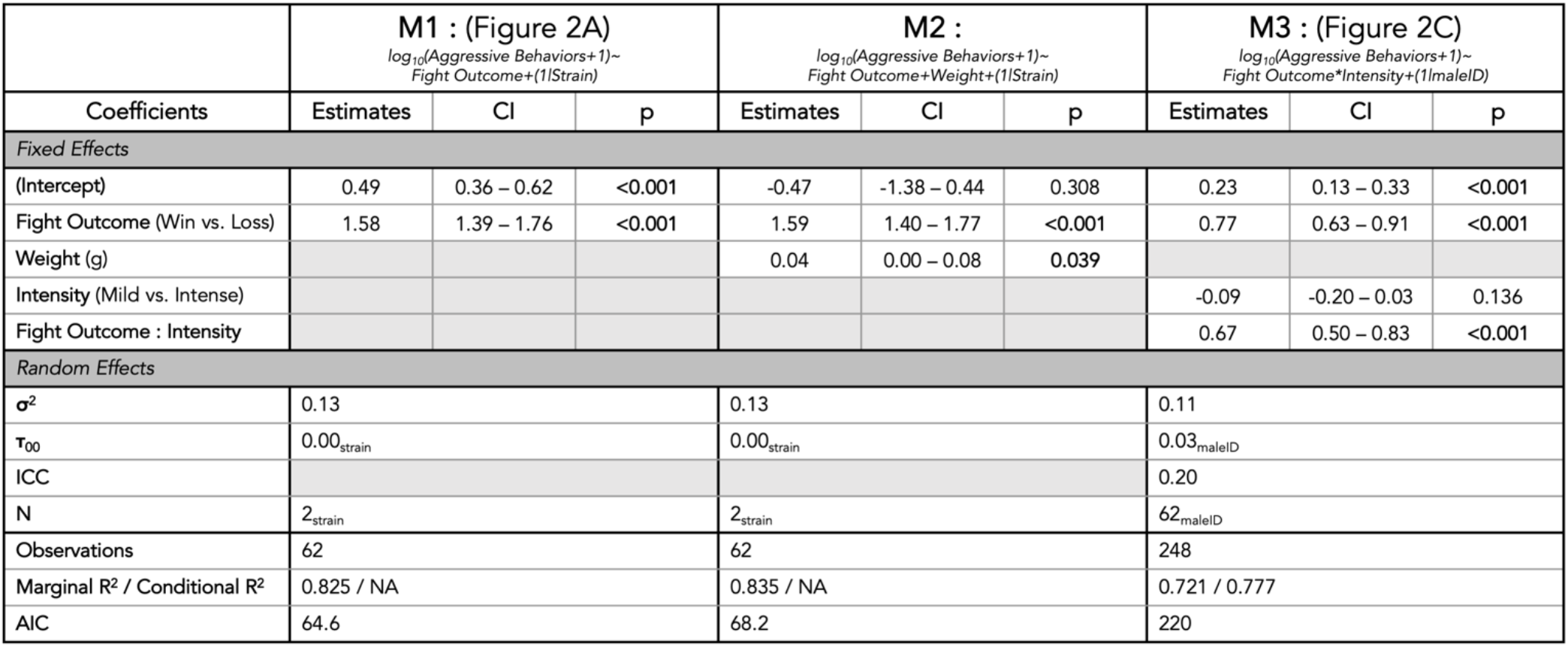
Linear mixed model (LMM) details accompanying results section 3a (Figure 2). The response variable for all three models (M1-M3) is the number of aggressive behaviors (count), and is logarithmically transformed in all cases.

**Table S2.**
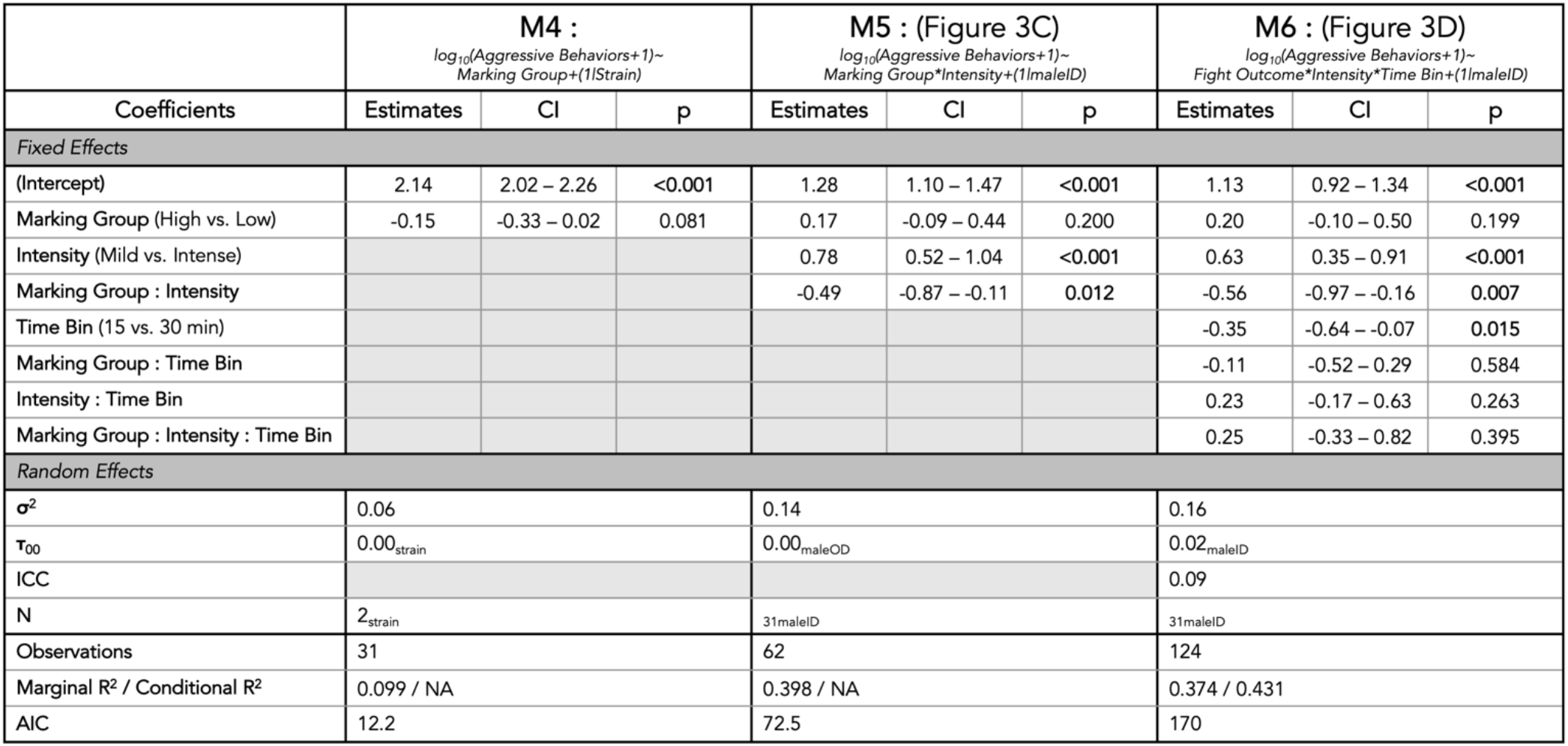
Linear mixed model (LMM) details accompanying results section 3b (Figure 3). The response variable for all three models (M4-M6) is the number of aggressive behaviors (count), and is logarithmically transformed in all cases.

**Table S3.**
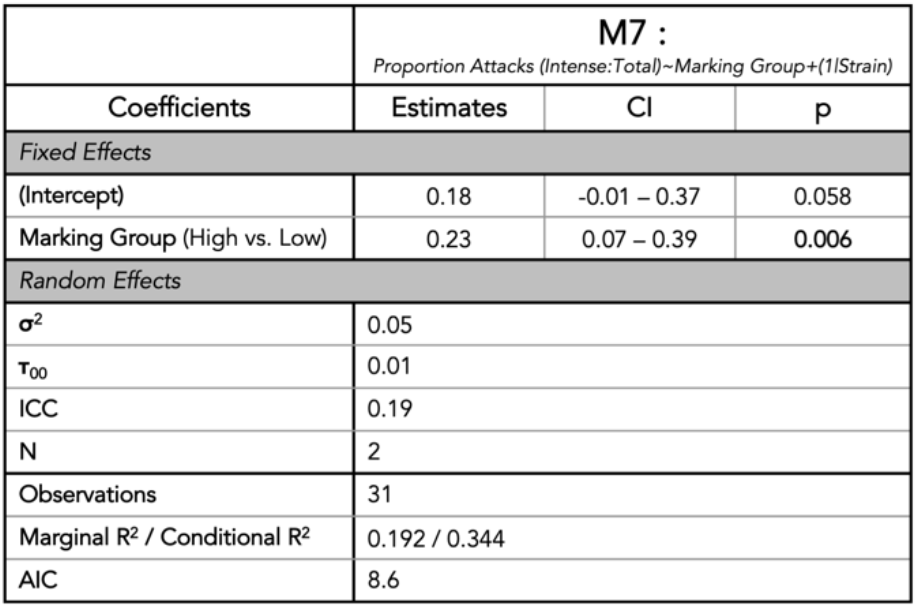
Linear mixed model (LMM) details accompanying results section 3c (Figure 4). The response variable is the proportion of intense : total aggressive behaviors.

|  | M7 :<br>*Proportion Attacks (Intense:Total)~Marking Group+(1\|Strain)* |  |  |
| --- | --- | --- | --- |
| Coefficients | Estimates | CI | p |
| **Fixed Effects** |  |  |  |
| (Intercept) | 0.18 | -0.01 – 0.37 | 0.058 |
| Marking Group (High vs. Low) | 0.23 | 0.07 – 0.39 | 0.006 |
| **Random Effects** |  |  |  |
| $\sigma^2$ | 0.05 | | |
| $\tau_{00}$ | 0.01 | | |
| ICC | 0.19 |  |  |
| N | 2 |  |  |
| Observations | 31 |  |  |
| Marginal R<sup>2</sup> / Conditional R<sup>2</sup> | 0.192 / 0.344 |  |  |
| AIC | 8.6 |  |  |

**Table S4.** Linear mixed model (LMM) details accompanying Figure S4. The response variable for all models (M8-M10) is the number of aggressive behaviors (count) performed by losers (M8,M9) or winners (M10), and is logarithmically transformed in all cases.

| | M8 : (Figure S4A)<br>loser aggression ~ loser signaling<br>$\log_{10}(\text{Aggressive Behaviors}+1) \sim \text{Marking Group} * \text{Intensity} + (1\vert \text{strain})$ | | | M9 : (Figure S4A)<br>loser aggression ~ winner signaling<br>$\log_{10}(\text{Aggressive Behaviors}+1) \sim \text{Marking Group} * \text{Intensity} + (1\vert \text{maleID})$ | | | M10 : (Figure S4B)<br>winner aggression ~ loser signaling<br>$\log_{10}(\text{Aggressive Behaviors}+1) \sim \text{Fight Outcome} * \text{Intensity} * \text{Time Bin} + (1\vert \text{maleID})$ | | |
| --- | --- | --- | --- | --- | --- | --- | --- | --- | --- |
| Coefficients | Estimates | CI | p | Estimates | CI | p | Estimates | CI | p |
| **Fixed Effects** |  |  |  |  |  |  |  |  |  |
| (Intercept) | 0.37 | 0.17 – 0.56 | <0.001 | 0.39 | 0.21 – 0.58 | <0.001 | 2.12 | 2.00 – 2.24 | <0.001 |
| Marking Group (High vs. Low) | 0.02 | -0.25 – 0.29 | 0.895 | -0.04 | -0.30 – 0.23 | 0.792 | -0.06 | -0.23 – 0.10 | 0.454 |
| Intensity (Mild vs. Intense) | -0.46 – 0.09 | 0.191 | -0.06 | -0.32 – 0.21 | 0.672 | 0.00 | -0.17 – 0.17 | 1.000 | -0.46 – 0.09 |
| Marking Group : Intensity | 0.11 | -0.27 – 0.49 | 0.562 | -0.14 | -0.52 – 0.24 | 0.463 | -0.00 | -0.23 – 0.23 | 1.000 |
| **Random Effects** |  |  |  |  |  |  |  |  |  |
| $\sigma^2$ | 0.14 | | | 0.14 | | | 0.05 | | |
| $\tau_{00}$ | 0.00<sub>strain</sub> | | | 0.00<sub>strain</sub> | | | 0.00<sub>strain</sub> | | |
| N | 2<sub>strain</sub> |  |  | 2<sub>strain</sub> |  |  | 2<sub>strain</sub> |  |  |
| Observations | 62 |  |  | 62 |  |  | 62 |  |  |
| Marginal R<sup>2</sup> / Conditional R<sup>2</sup> | 0.041 / NA |  |  | 0.054 / NA |  |  | 0.018 / NA |  |  |
| AIC | 73.8 |  |  | 73.0 |  |  | 17.1 |  |  |

**Figure S1.**
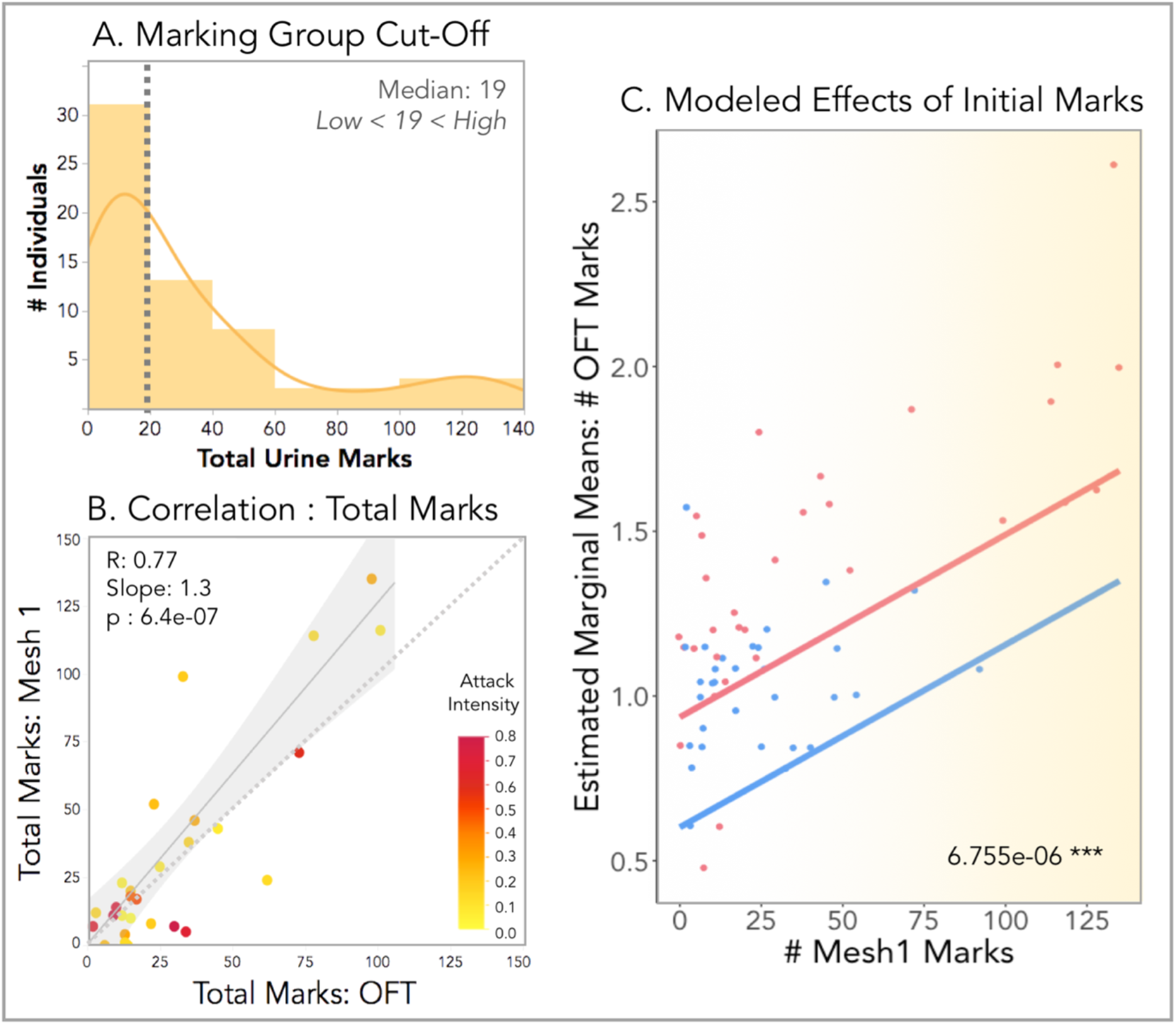
**(A)** Histogram of the distribution of total urine marks deposited by all males competitors in the Mesh Trial (pre-fight). The median (19) number of marks is indicated with dashed line. High-marking versus low-marking males were categorized based on whether the total marks deposited was either above or below the median. **(B)** Correlation plot of the total number of marks deposited by each winning male prior to the contest trial (Mesh 1) and 1 day after the contest in the open field trial (OFT) (48). The number of marks deposited pre- and post-fight are quite well correlated with each other (R=0.77). This is despite the differences in aggressive experience, arena size, and social stimulus in the environment across these two trials. Individual data points are color-scaled by the proportion of total attacks that were intense (red: high; yellow: low). The males that marked lowly in both trials (clustered in the bottom left corner) tend to perform more intense attacks (more red). One male was removed from this correlation analysis as an outlier (NY3-131), though excluding or including this male does not affect the overall pattern. **(C)** Estimated marginal means of the total number of OFT marks (log-transformed) given fight outcome (winner=red; loser=blue) and initial signal investment (# Mesh 1 marks). Initial signal investment significantly predicts marking levels of winners and losers post-fight. The model p-value of the effect of the # Mesh 1 marks on the total OFT marks indicated in the bottom righthand corner (highly significant).

**Figure S2.**
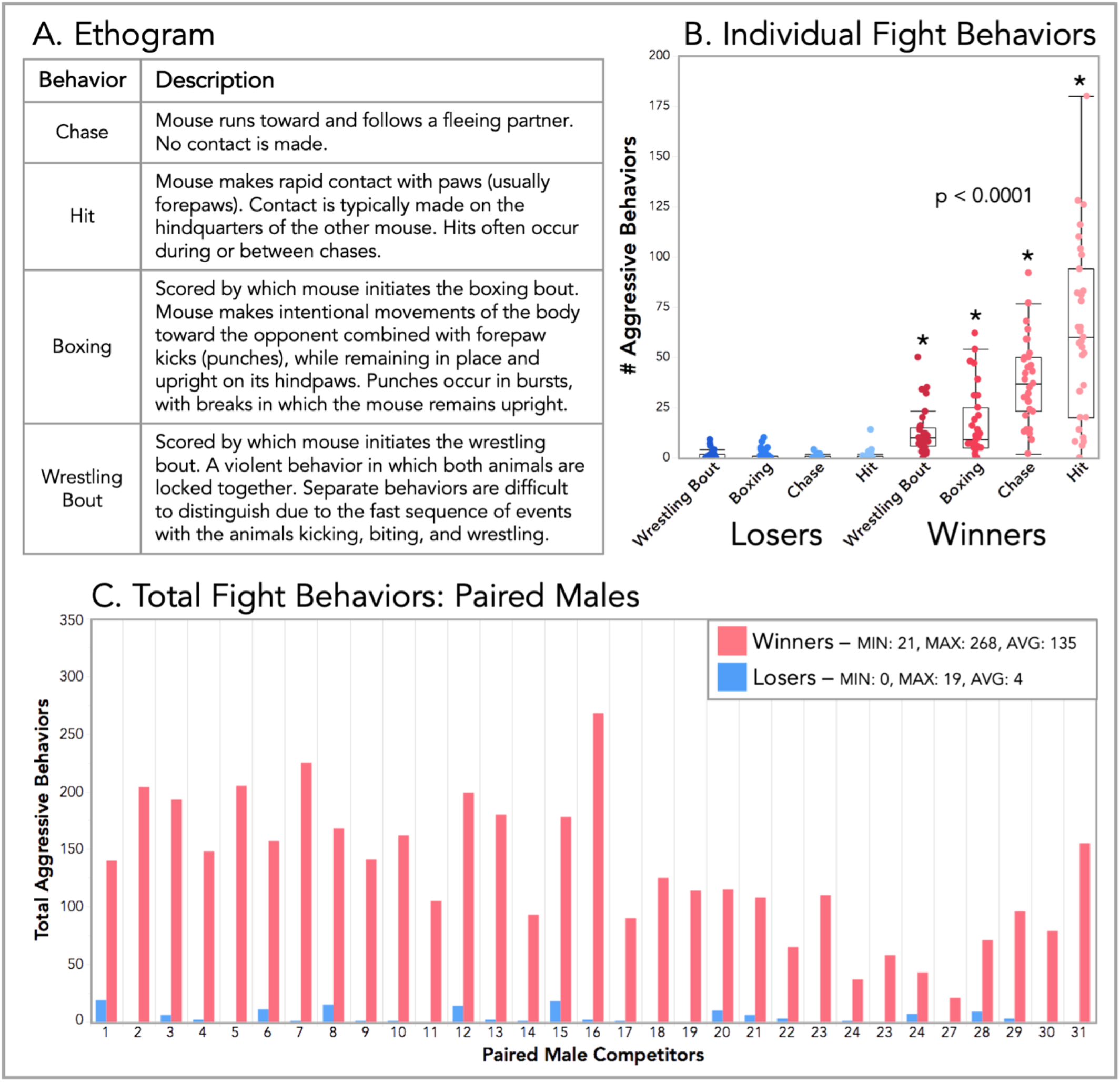
Fight trial behaviors. **(A)** Ethogram of scored aggressive behaviors. **(B)** Boxplots of the total counts for individual fight behaviors performed by winning (red) and losing males (blue). Winners performed significantly higher levels of each aggressive behavior compared to losers (* p<0.0001) **(C)** Total aggressive behaviors performed by each paired male competitor (31 pairs). Winners indicated in red and losers in blue. The fight outcome (the categorization of winners and losers) was determined by which male performed more aggressive behaviors within a pair. Across all pairs, winners ranged from performing 21-268 attacks, and losers ranged from performing 0-19 attacks.

**Figure S3.**
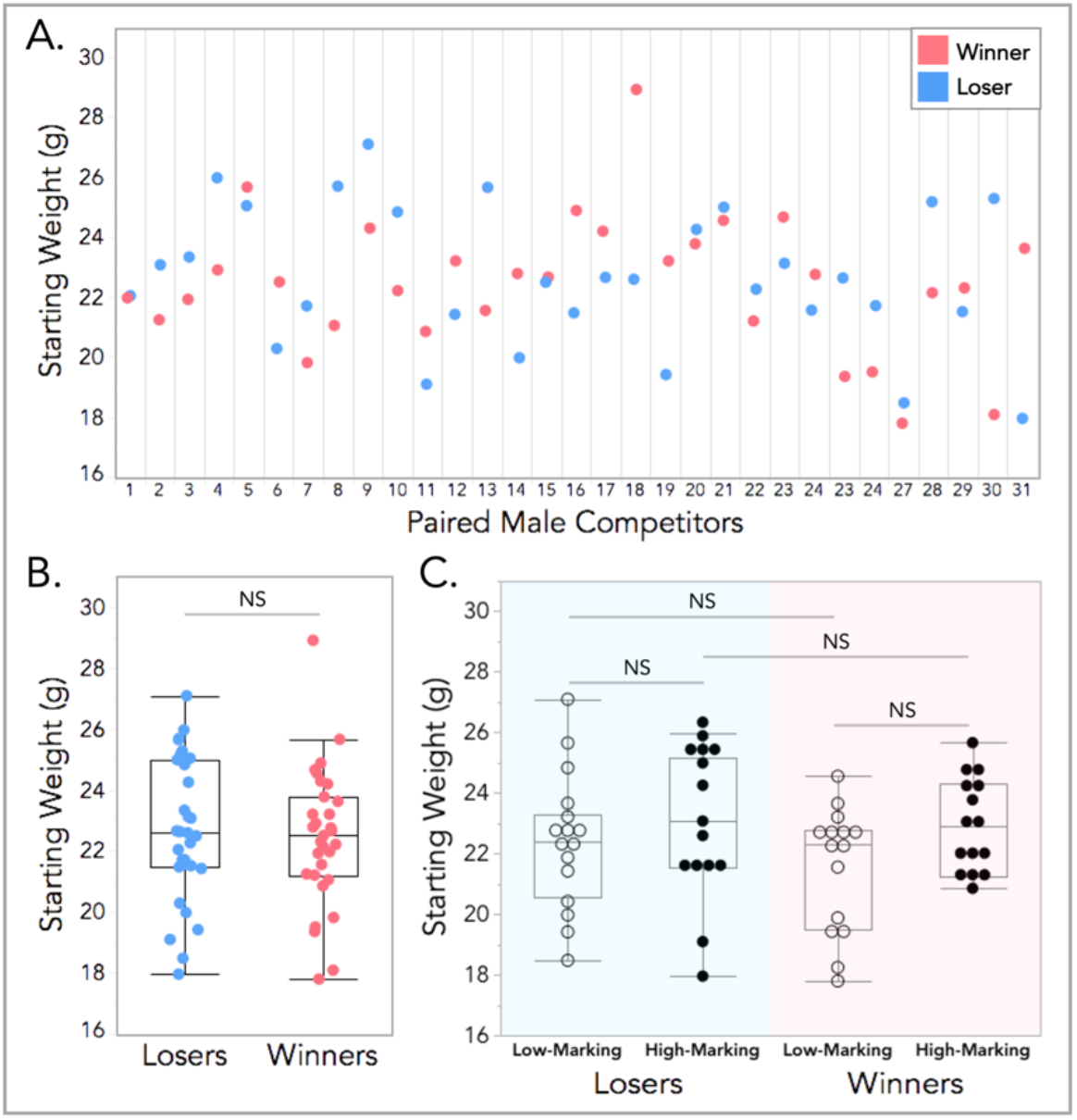
Body weight differences across winning and losing males. All weight measurements (g) were collected the day before trials began, such that competitors were weight-matched as closely as possible. **(A)** Starting weights (g) of each male within a competing pair across trials. **(B)** Boxplot of starting body weights of all losers and winners. Winners and losers did not differ in weight across trials. **(C)** Boxplot of the starting body weight for winners and losers by marking group: high vs. low-marking males. A single male was removed from this analysis as an outlier (NY2-205), however removal of this male did not affect the observed patterns. A LMM was used to model relationships, and analyses of variance were used to test for overall effects. Significance codes: NS p>0.05.

**Figure S4.**
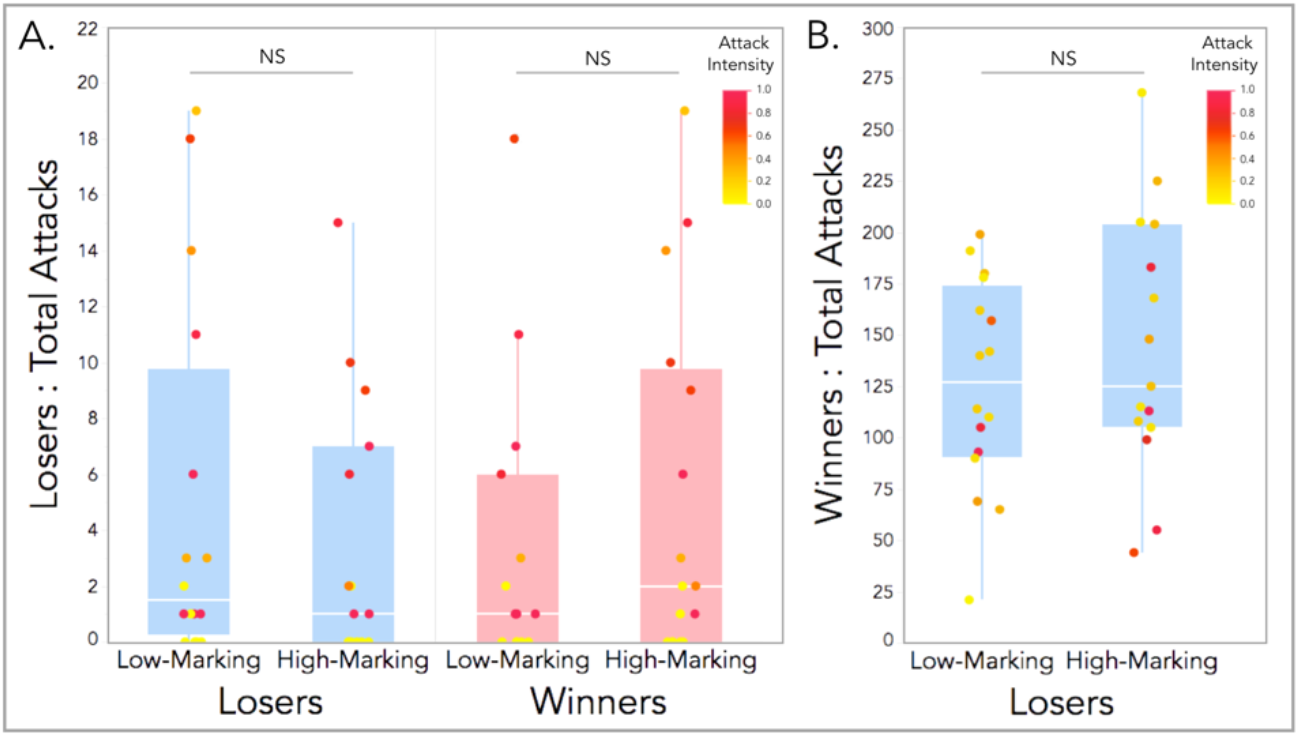
Additional dyadic comparisons of contest aggression and intensity in response to signaling effort. **(A)** Boxplot of the total attacks performed by losers given either loser or winner marking investment (low vs. high). The initial signaling effort of losers or winners does not predict loser aggression, both in terms of total attacks or intensity. Though losers perform very few attacks and only early on in the trial. **(B)** The initial signaling effort of losers similarly does not predict winner aggression, in total attacks or intensity. LMMs were used to model relationships, and analyses of variance were used to test for overall effects. Significance codes: NS p>0.05.

## Ethics

All experimental protocols conducted at Cornell University were approved by the Institutional Animal Care and Use Committee (IACUC: Protocol #2015-0060) and were in compliance with the NIH Guide for Care and Use of Animals.

## Data accessibility

Data sheets and R code used in all analyses are available on the Dryad Digital Repository.

## Authors’ contributions

C.H.M and M.J.S. conceived the study. C.H.M. performed trials and analyses. M.F.H and K.H. scored behavioral trials. T.R. created figure content. All authors contributed to manuscript preparation.

## Competing interests

The authors declare no competing interests.

## Funding

This research was funded by USDA Hatch Grant (NYC-191428; Michael Sheehan) The funders were not involved in the design of the study; the collection, analysis and interpretation of data; the writing of the manuscript and any decision concerning the publication of the paper.

## Acknowledgements

We thank Kevin Besler, Kusuma Anand, Christen Rivera-Erick and Melanie Colvin for crucial technical assistance; Russell Ligon and Caleb Vogt for helping establish recording systems and tracking methods in the lab; and James Tumulty for manuscript feedback.

## Notes

### Competing Interest Statement

The authors have declared no competing interest.

